# Spatiotemporal regulation of cell cycle states within the complex tumor microenvironment

**DOI:** 10.64898/2025.11.28.691200

**Authors:** Gianni Zanardelli, Olivier Tassy, Maulik K. Nariya, Nacho Molina

## Abstract

Tumor growth and resistance arise from the interplay between cell-cycle dysregulation and the spatial organization of the tumor microenvironment (TME). While spatial transcriptomics now enables molecular profiling of intact tissues, it captures only static molecular states, making it challenging to reconstruct dynamic processes such as proliferation. Here, we develop SpaceCycle, a computational framework that infers the continuous cell-cycle phase and associated oscillatory gene expression dynamics from spatial transcriptomic data. Using human melanoma, breast, and lung tumors, we map both discrete and continuous cell-cycle states across entire tissue sections to reveal how proliferative activity is spatially structured within the TME. We find that cycling and non-cycling cells form distinct spatial niches reflecting differences in vascularization, immune composition, and cellular density. Continuous-phase inference further exposes tumor-specific variations in cell cycle phase durations and uncovers oscillatory programs. Together, our results provide a spatiotemporal view of cell-cycle organization in human tumors and establish a general framework for detecting dynamic transcriptional programs and spatial proliferation patterns from static tissue data.

## Introduction

Cancer is a group of complex diseases in which cells acquire deleterious mutations, particularly in genes that regulate the cell cycle, suppress abnormal growth, or repair DNA damage. Genetic alterations are necessary for cancer initiation, however, they are not sufficient to promote tumor growth and metastasis. It has long been recognized that cancer cells reside within a complex ecosystem called the tumor microenvironment (TME), which includes diverse components such as stromal and immune cells, blood vessels, the extracellular matrix (ECM), and signaling molecules like growth factors, cytokines, and chemokines. Over the past few decades, it has become abundantly clear that the behavior of cancer cells, in particular how they grow, invade and resist therapy, depends strongly on the TME. Early discoveries showed that tumors require new blood vessels to grow beyond a certain size [1–3], by the 2000s, it was established that stromal cells such as fibroblasts and macrophages, as well as the ECM, actively support tumor progression rather than being passive bystanders [4–6] and more recently, immune checkpoint blockade discoveries revealed how tumors can suppress anti-tumor immunity, leading to therapies that have transformed cancer treatment [7–9].

The advent of single-cell and spatial omics technologies has now enabled an unprecedented view of tumor organization. Spatial transcriptomics and multiplexed imaging methods such as MERFISH, CyCIF, and GeoMX allow quantification of hundreds to thousands of transcripts or proteins across entire tissue sections while preserving spatial context [10–12]. This has reinvigorated efforts to map the cellular composition, interactions, and metabolic gradients within tumors at subcellular resolution. Yet, despite these advances, certain fundamental biological processes remain unexplored in situ.

One such unexplored area in histopathology of cancer is tissue-wide identification of the cell cycle states. The cell cycle is a highly regulated biological process during which cells go through the cell cycle phases–G1, S, G2, and M, and divide into two daughter cells; the cells can also temporarily exit the cell cycle and enter a quiescent state, often referred to as G0 phase, while maintaining their ability to re-enter the cell cycle, typically in the G1 phase. It is well established that the cell cycle is highly dysregulated in cancer and is often characterized by loss of checkpoint control at G1/S and S/G2 transitions, overexpression of cyclins and cyclin-dependent kinases (CDKs), accumulation of DNA damage, replication stress, and genomic instability, making it a promising target for the development of new therapeutic strategies. While much is known about cell-cycle phase transitions in isolated cells in culture or in bulk populations, far less is understood about how cells in situ within tissues coordinate their proliferation with neighboring cells and how this organization manifests spatially within tumors. The TME is characterized by nutrient gradients, hypoxia, variations in cellular density, and heterogeneous immune infiltration, but how specific cell-cycle states correlate with these microenvironmental features remains poorly understood. For example, it is not well understood whether proliferating cells preferentially localize near the vasculature or whether quiescent cells dominate hypoxic regions. Hypoxia is known to push cells into quiescence, and quiescent states are increasingly recognized as important drivers of therapy resistance and immune evasion in cancer. Identifying where quiescent cells reside in the tissue may therefore have direct clinical value [13, 14]. It is also not known whether the cellular densities are uniform across cell cycle phases or whether there are spatial niches based on cell cycle states, i.e. do the cells tend to surround themselves by other cells that are in the same cell cycle phase? Mapping cell cycle states across tissues could bridge the gap between molecular and histological perspectives of tumor biology, offering new biomarkers for tumor aggressiveness and treatment response.

Addressing these questions requires integrating spatial context with temporal information about the cell cycle. Recent advances in computational biology have made it possible to infer continuous cell-cycle progression from transcriptomic data. Methods that incorporate RNA velocity and biophysical modeling, such as DeepCycle and FourierCycle, can reconstruct oscillatory transcriptional programs underlying cell proliferation [15–18]. However, such dynamic modeling has so far been restricted to dissociated single-cell sequencing datasets, which lack spatial resolution.

In this study, we integrate spatial transcriptomics with cell cycle inference to map proliferative dynamics within intact human tumors. Using publicly available spatial omics data from melanoma, breast, and lung cancers, we identify the major cellular populations within each tumor, estimate their discrete and continuous cell cycle states, and relate these states to their spatial organization. We show that proliferating and quiescent cells form distinct microenvironmental niches, reflecting local differences in vascularization, immune infiltration, and tissue density. Finally, we introduce SpaceCycle, an expectation–maximization framework for pseudotime inference that uncovers oscillatory gene expression programs that distinguish subpopulations of cancer cells within the TME. Together, our results provide a spatially resolved view of tumor cell-cycle dynamics and reveal new principles of proliferative organization in human cancers.

## Results

### The TME is rich with diverse cell types and complex cellular interactions

To investigate the influence of the TME on the cell cycle states we used publicly available spatial transcriptomics data of formalin-fixed paraffin-embedded (FFPE) tumor samples obtained from human patients [19]. The samples were profiled using a panel of 500 genes pertinent to human immuno-oncology and contained quantification of millions of mRNA transcripts across whole tissue sections (see Table S1). In this work, we focus on melanoma, breast, and lung tumors, but our methodology is general and can be applied to other samples. We used the Leiden algorithm to cluster the cells based on their transcriptional similarity and then we used cell markers to assign cell types to every cluster (see **Methods** for details) [20–22]. Fig. 1 shows cell clusters visualized in a UMAP embedding and the spatial map of the different cell types within the tumor.

**Figure 1.**
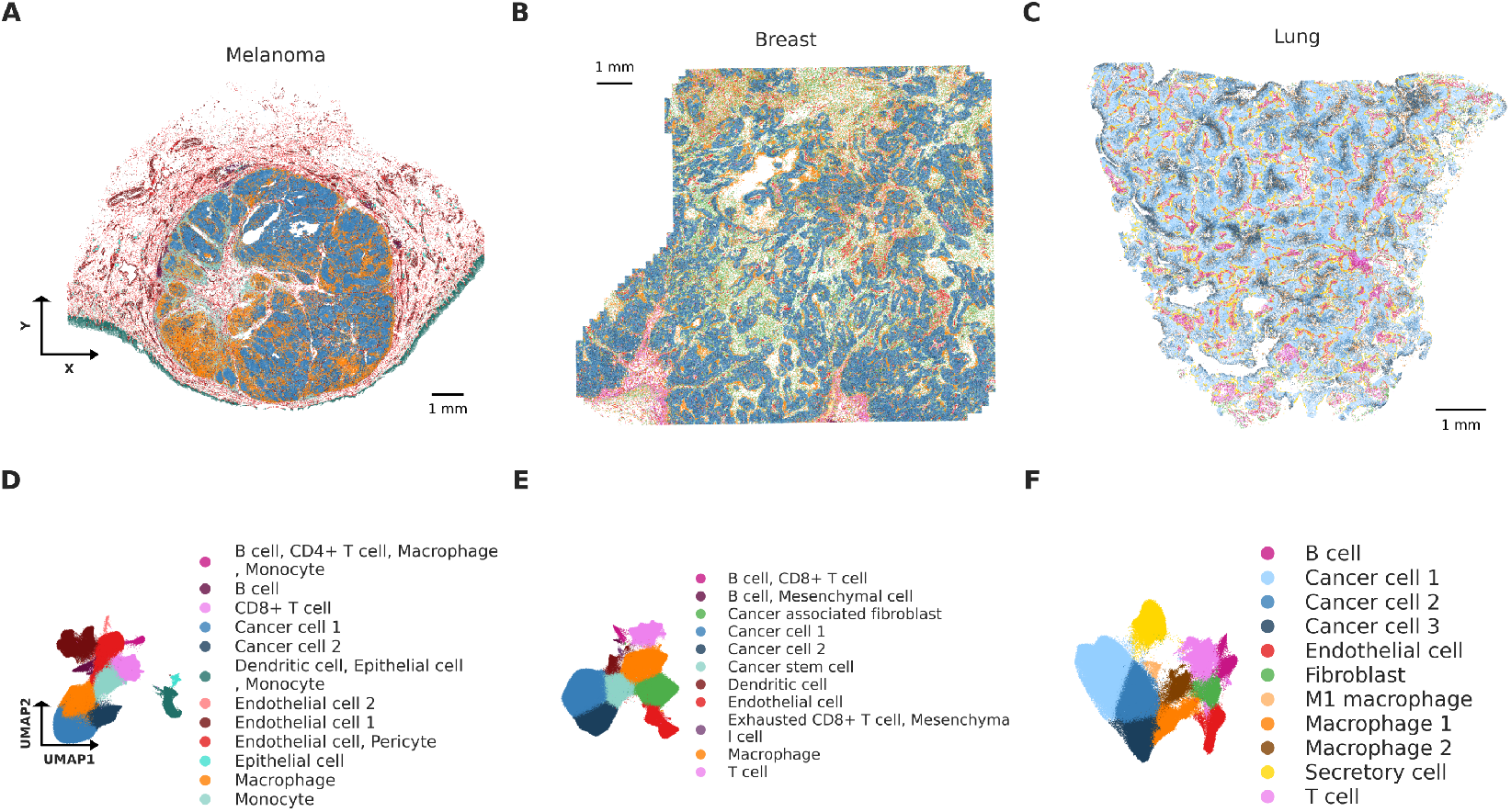
Spatial map of the cell types shows diversity in cell types across the three human tumors. **A-C:** Spatial map of the cell types within the melanoma, breast, and lung tumors respectively, based on the transcriptomic profiles. **D–F:** the corresponding UMAP embeddings of cells in the tumors. Clustering was performed using the Leiden algorithm, whereas cell type identification was done using the top 30 highly expressed tissue-specific gene markers of known cell types obtained from the CellMarker2.0 databse. The TME is comprised of diverse cell types and exhibits complex cellular interactions

In the melanoma tumor (see Fig. 1A and 1D), we found that among the distinct cell clusters, cancer cells of epithelial origin, macrophages, endothelial cells and monocytes were the most abundant cell types in the tumor (see Table S2). Monocytes and macrophages are critical regulators of the TME, influencing tumor progression, immune evasion, and therapeutic outcomes [23– 25]. A large number of endothelial cells and pericytes suggests that there is a high degree of vascularization which often reflects aggressive and rapidly growing tumor phenotype. In the breast tumor (see Fig. 1B and 1E), in addition to the cancerous luminal epithelial cells and immune-associated cells, we found an abundance of cancer-associated fibroblasts (CAFs) and cancer stem cells throughout the tumor. Both CAFs and cancer stem cells are critical for shaping the TME and supporting tumor growth through a variety of mechanisms— ECM remodeling, growth factor secretion, self-renewal, immunomodulation, supporting TME cellular niches, and metabolic reprogramming to name a few [6, 26, 27]. Interestingly, we found two sub-populations of the epithelial cells in the tissue referred to as cancer cell 1 in blue and cancer cell 2 in dark blue (both of these comprise of luminal epithelial cells) in Fig. 1E. We found that there was a distinct spatial patterning among these cells, in that the epithelial 2 cells tend to cluster together and tend to be surrounded by epithelial 1 cells; we found a similar and more prominent patterning in the lung tumor, indicative of functional compartmentalization and complex cellular interactions within the TME. Lastly, the lung sample (see Fig. 1C and 1F) was comprised of tissue specific cells such as basal cells, which are the principle stem cells of the airway epithelium and differentiate into all major cell types in lung, and secretory cells. Basal cells are the main cell of origin for squamous cell carcinoma and it is known that basal-like molecular signatures are associated with poor differentiation, higher aggressiveness, and resistance to therapy in non-small cell lung cancer [28–30]. In summary, the microenvironment of the samples was comprised of a diverse mixture of cells giving rise to complex cellular interactions with the TME. In the following sections we will uncover the effect of these interactions on the cell cycle states in the TME.

### Discrete cell phase identification reveals subtypes of cancer cells with distinct cell cycle state composition within the TME

In order to understand the effect of TME on the cell cycle, we identified the cell cycle phases of the constituent cells in the TME. We used known cell cycle phase markers to assign G0, G1, S, and G2/M scores to every cell in the tumor. To improve the interpretability of our results we used a softmax function to convert the cell cycle scores into probabilities. This approach allows us to filter the cells on the basis of the certainty we have on their phase assignment (see details in the **Methods** section and Fig. S1).

Fig. 2A, 2B, and 2C show the spatial distribution of cell cycle phases—G1 in blue, S in red, G2/M in green and G0 in yellow—for melanoma, breast and lung samples respectively (Supplementary Fig. S2 shows the spatial distribution of discrete cell cycle phases of only the cancer cells in the tumor samples). We found, by visual inspection, that the cycling cells and the quiescent cells tend to clump among themselves, suggesting that there are metabolically active and inactive regions within the TME. We quantify this by calculating the local density of quiescent (non-cycling) and proliferating (cycling) cells across the tissue in the proceeding section. Fig. 2D, 2E, and 2F show the proportions of cells in discrete cell cycle phases, including G0 cells, for each cluster in the melanoma, breast and lung tumor respectively. We found that endothelial cells were among the most abundant quiescent cells in all the three tumors. Quiescent endothelial cells are known to suppress cancer invasiveness and proliferation, while reprogrammed and proliferative endothelial cells promote angiogenesis [31, 32]. Following endothelial cells, the immune populations such as B cells, T cells, macrophages, and monocytes had the highest proportions of non-proliferating cells across the three tumors. Lastly, the breast sample had a significant proportion (96.9%) of cancer-associated fibroblasts (CAF) and the lung sample had a significant proportion (98.9%) of fibroblasts that were identified to be in G0 phase. Quiescent fibroblasts eventually evolve into CAFs which remodel the ECM, promote angiogenesis, and support tumor growth; their initial quiescent state is thought to be tumor-restraining, however, activation of these cells is a key step involved in creating a pro-tumorigenic environment [33–35]. In conclusion, we obtained a spatial map of discrete cell cycle states across whole tissue sections of tumor samples, as well as quantified cell phase proportions of the constituent cells in the TME. Our approach addresses a previously unexplored aspect of tumor spatial organization by linking cell cycle states with the tissue architecture at single-cell resolution.

**Figure 2.**
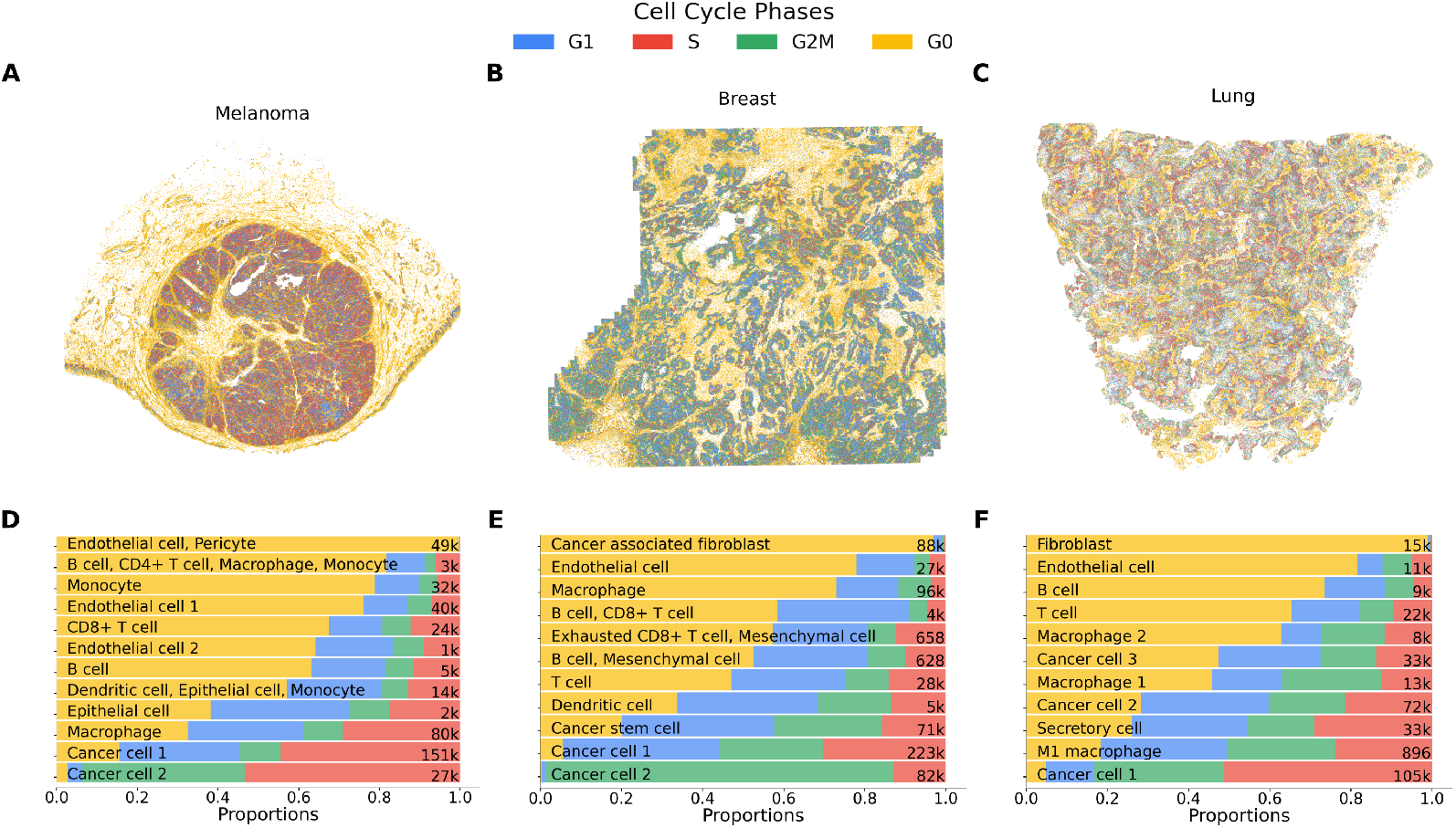
Identification of discrete cell cycle phases of the cells in the TME. **A-C:** show the spatial localization of the cell cycle phases in the melanoma, breast, and lung tumors respectively, the cells are colored according to their assigned phase: G1 in blue, S in red, G2/M in green, and G0 (quiescent) in yellow. Cell cycle phases were identified by calculating G0, G1, S and G2/M score based on the expression of cell cycle marker, the scores were then transformed into probabilities using a softmax function. **D-F:** show proportions of cells in G0, G1, S, and G2/M phases across the cell clusters in melanoma, breast and lung respectively. Distinct proportions of cycling states are observed among malignant cell types, highlighting differences in proliferative capacity among cancer cells in the TME.

### Cycling and non-cycling cells form a cellular niche within the TME

To further investigate the association between the cell cycle states and the tissue architecture we examined the cellular densities in the local neighborhood of the TME. In Fig. 3A-C, we visualize the cellular densities of all cells across the melanoma, breast and lung samples respectively. We defined the local neighborhood of a cell as a region with a radius of 50 micrometers surrounding the cell, which is roughly 2–5 cell diameters for most mammalian epithelial cells. In supplementary Fig. S3, we visualize the cellular densities cancer, immune and stromal compartments separately. We found that there was a large variability, close to two orders of magnitude, in the cellular densities across sections of the tumor samples, which is indicative of strong spatial heterogeneity in tumor organization and function. It is known that high cellular density is indicative of rapid proliferation, poor vascularization, and restricted diffusion of nutrients and drugs, whereas low cellular density regions mark necrotic areas, stromal regions, or immune cell infiltration zones [36–38].

**Figure 3.**
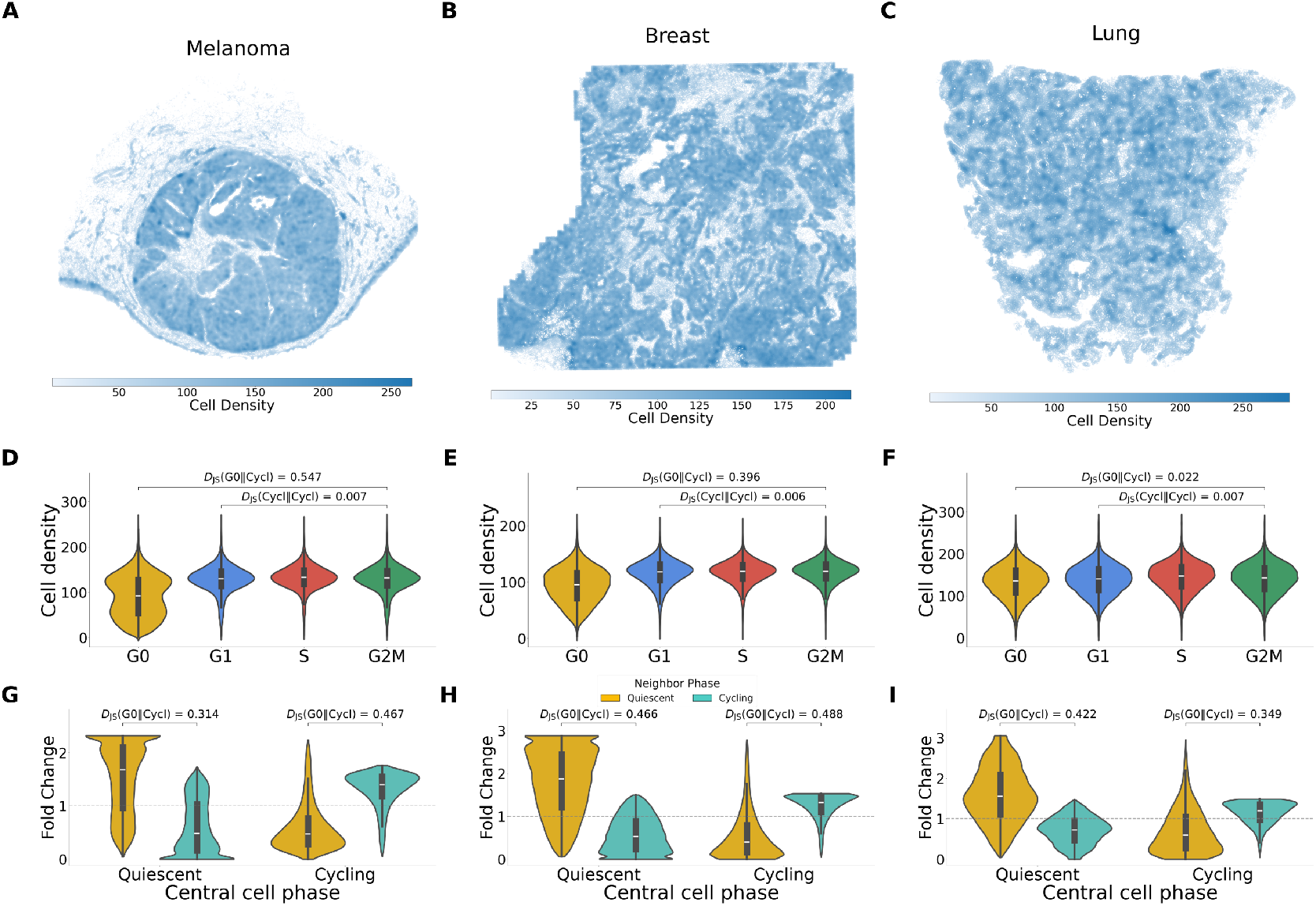
Densities of cycling and non-cycling cells reveal cellular niches in the TME. **A-C**, Cellular density in the local neighborhood, within 50 micron radius, in melanoma, breast, and lung sample respectively. **D-F**, Violin plots showing distribution of cellular densities stratified by the cell cycle state of each observed cell in the melanoma, breast, and lung tumors respectively. Quiescent cells show high variability in their cellular density and tend to localize in low density regions compared to cells that are in G1, S or G2/M phases, note that we used the Jensen-Shannon divergence to quantify the differences in the distributions (see **Methods**) **G–I**, Violin plots showing the enrichment in the neighborhood cell state composition based on the cycling state of the observed cell in the three tumors, cycling in blue, non-cycling in yellow. Non-cycling cells tend to localize in regions with an abundance of non-cycling cells whereas cycling cells tend to localize in regions with an abundance of cycling cells, indicating presence of non-proliferative and proliferative cellular niches in the TME.

To add a new dimension to the cellular density analysis we grouped the cells according to their discrete cell cycle phases and calculated the densities for each group. As shown in Fig. 3D-E, we found that the distribution of the cellular densities of the “cycling cells”, i.e. cells that are either in G1, S or G2/M phases were similar to one another across the three samples. Interestingly, we found a large variability in the cellular densities of “non-cycling” cells, i.e. cells that were identified as being in G0 phase, especially in the melanoma and the breast tumor. Lastly, we grouped the cells into “non-cycling” and “cycling” and calculated the cellular densities of each group with respect to itself and with respect to the other.

Strikingly, we found that the non-cycling cells show enrichment of other non-cycling cells in their local neighborhood, whereas the cycling cells show an enrichment of other cycling cells in their local neighborhood. This trend was consistent across all three tumor samples, see Fig. 3G-I, indicating that the cycling and non-cycling form cellular niches in the TME. To the best of our knowledge, such a grouping by cell cycle states has not been previously reported in the tissue architecture of tumors.

### SpaceCycle reveals oscillatory gene expression dynamics of cancer cells within the TME

Lastly, we performed pseudotime analysis, a computational technique to order cells based on their gene expression profiles, to infer the continuous cell cycle phase of cancer cells in the TME. We modeled the gene expression using a Fourier series and used an expectation-maximization (EM) algorithm (details in **Methods** section) to infer the cell cycle phase as well as the expression dynamics. Briefly, the EM algorithm is an iterative method to determine a local maximum likelihood estimate of the parameters of the model, in this case the Fourier coefficients of the gene expression dynamics, where the model depends on a latent or unobserved variable, in this case the continuous cell cycle phase, *θ*, of the cell. In order to validate this approach, we used SpaceCycle to obtain continuous cell cycle phase *θ* from single-cell RNA-seq profiles of mouse embryonic stem cells (mESCs) and human fibroblasts. As shown in supplementary Fig. S4, we found an excellent agreement between *θ* as well as the phase transition boundaries between SpaceCycle and DeepCycle [16]. We used this approach to infer the continuous cell cycle phase for the subpopulation of cancer cells in each tumor sample and obtained the oscillatory expression dynamics of gene markers for these cells.

Fig. S5 shows the proportions of cells in G1, S and G2/M phases with respect to the continuous cell cycle phase, *θ*, for the subpopulations of cancer cells in melanoma, breast, and lung. When aggregated, the proportions concur with the proportions found with discrete phase identification reported in Fig. 2. However, with the continuous phase identification, we can now determine the relative cell phase durations for the cancer cells. This allows us to estimate the fraction of time that a given population of cells spends each phase relative to total cell cycle duration. For instance, in the melanoma specimen we found that the cells in cancer cell 1 cluster (see supplementary Fig. S5A) spent 31% of time in G1 and 69% of the time in S and G2/M was extremely short-lived, suggesting accelerated cell cycle progression due to loss of DNA damage checkpoints, whereas the cells in cancer cell 2 cluster (see supplementary Fig. S5D) spent 54% of time in S and 46% of the time in G2/M and the G1 phase is extremely short-lived indicating unrestricted cell cycle entry. In contrast, in the breast specimen, cancer cell in cluster 1, Fig. S5B, spent roughly equal times in the G1, S, and G2/M phases, whereas cells in cancer cell 2 cluster, Fig. S5E, spent 95% time in G2/M, suggesting G2 arrest in response to DNA damage. Lastly, in the lung specimen, cells in cancer cell 2 spent roughly equal times in the cell cycle phases, whereas cells in cancer cluster 1 have a short G1 phase, once again indicating checkpoint loss and uncontrolled proliferation (see supplementary Fig. S5C and S5F).

In addition to determining the relative time spent in the cell cycle phases, SpaceCycle allows to inspect the gene expression dynamics in the cancer subpopulations. Fig. 4 shows the gene expression dynamics of top 30 genes exhibiting an oscillatory dynamics during the cell cycle in the cancer cells. We show oscillatory dynamics of markers of mitosis, such as Aurora Kinase A and B (AURKA and AURKB) across the tumor samples. AURKA and AURKB promote spindle assembly and maintain the spindle assembly checkpoint during the prometaphase, both AURKA and AURKB are important to preserve the malignant state [39–42]. We also found the oscillatory dynamics of markers of S-phase, such as the MutS homologs 2 and 6 (MSH2 and MSH6), notably known for their role in the mismatch repair mechanism [43]. In addition to known regulators of the cell cycle. we also found genes that are not directly related to the cell cycle but appear to show an oscillating behavior, such as EPHB3 in the population of malignant cells in the breast tumor or FBLN1 in the population of lung cancer cells. Ephrin type-B receptor 3, or EPHB3, can participate in both inhibition [44] and proliferation [45] of cells, and in cancer, it can be a tumor repressor [46] while its knockdown can also be tumor suppressive [47], which makes it an ambivalent gene for both oncology and cell cycle. Fibulin-1, in non-small cell lung cancer, has isoforms C and D that acts as EGFR suppressor [48], EGFR being upregulated in lung cancer participates to the tumor progression [49].

In summary, our pseudotime analysis allows us to obtain a mechanistic view of the differences in cell proliferation of the subpopulations of cancer cells in the TME and reveals previously unrecognized patterns in gene expression dynamics of regulators of tumor progression.

**Figure 4.**
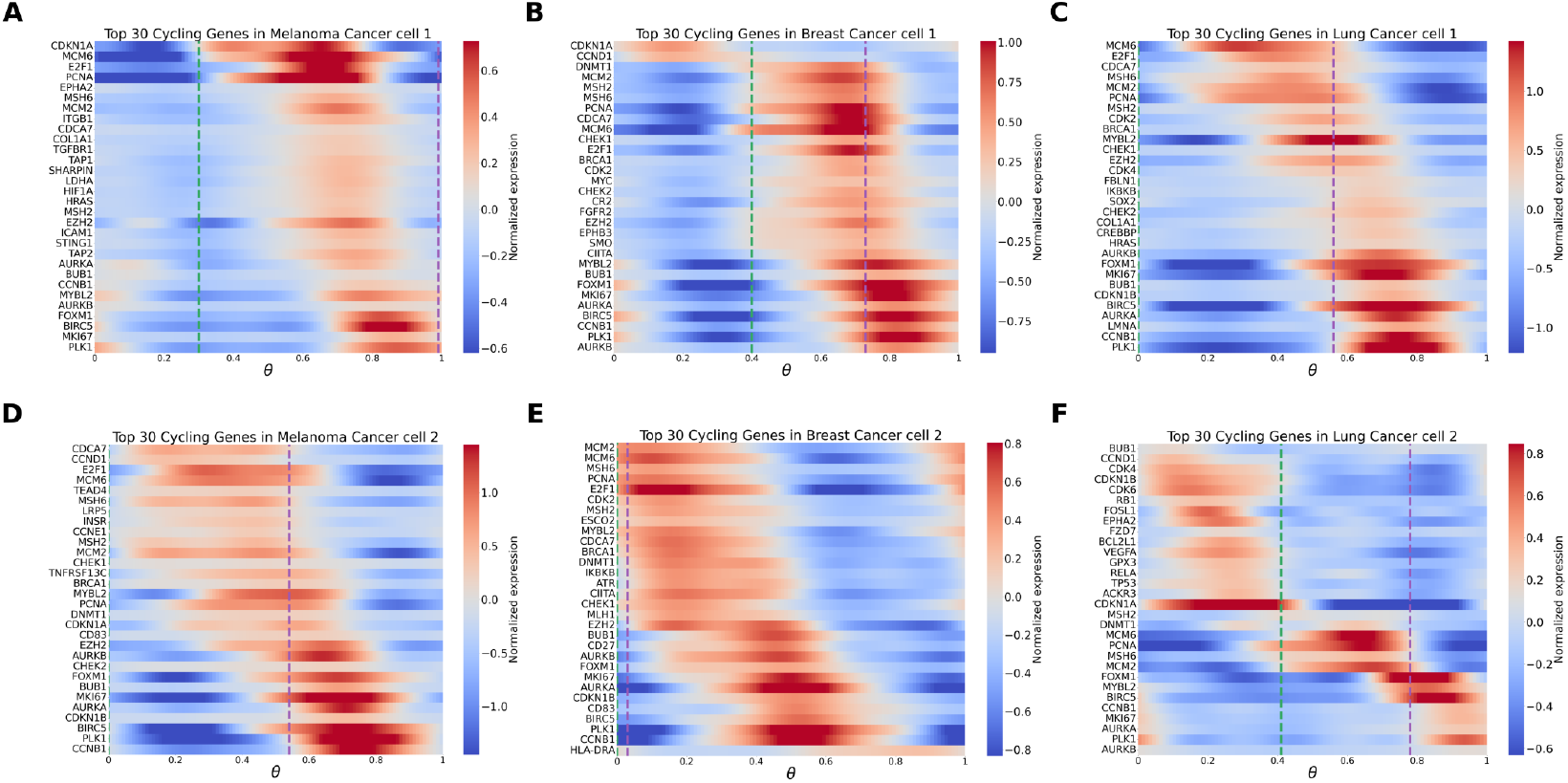
SpaceCycle enables identification of continuous cell cycle phase, gene expression dynamics, and reveals distinct proliferative dynamics among the cancer cells in the TME. Gene expression dynamics of top 30 cycling genes versus the continuous cell cycle phase identified by SpaceCycle for the cell clusters in the tumors, melanoma cancer cell 1 in **A** and cancer cell 2 in **D**, breast cancer cell 1 in **B** and cancer cell 2 in **E**, and lung cancer cell 1 in **C** and lung cancer cell 2 in **F**

### Limitations of the study

The spatial datasets analyzed here provide high spatial resolution but are based on targeted MERFISH panels acquired from formalin-fixed, paraffin-embedded tissue. As a result, only a subset of pathways and regulators involved in cell-cycle control, quiescence, and immunity is captured, and RNA degradation may affect the detection of cell cycle-associated transcripts. Consequently, our inferences about regulatory programs and microenvironmental cues are necessarily restricted to the genes present on the panel and those that remain robustly detectable in FFPE samples.

All cell-cycle and quiescent states in this study are inferred from transcriptomic profiles alone, without coregistered in situ ground truth such as protein levels of Ki-67, FUCCI reporters, or EdU/BrdU labeling in the same samples. While we use established gene sets and filtering strategies, quiescence is particularly challenging to define from RNA data and may in part reflect the absence of cycling markers rather than a fully characterized G0 program.

Within the SpaceCycle framework, pseudotime inference is performed separately for each cell cluster, so the inferred cell-cycle progression (ranging from 0 to 1) represents a *relative* timescale that is specific to each population. This means that the same pseudotime interval may correspond to different real-time cell-cycle durations across clusters or tumor types, and our interpretations of “short” or “long” phases should be viewed as comparative rather than absolute. Moreover, the underlying model assumes that cyclic dynamics can be described by a low-order Fourier basis and that cells are sampled from a quasi–steady-state distribution along the cycle, which may not fully capture more complex or non-periodic behaviors.

Our spatial analyses are based on a single two-dimensional section from one tumor per cancer type. This design allows us to illustrate the feasibility and biological relevance of the approach, but it does not capture three-dimensional tissue architecture or inter-patient variability, and therefore should be regarded as a proof of concept rather than a population-level characterization. Finally, SpaceCycle infers oscillatory dynamics from static snapshots; direct validation of these inferred trajectories and phase durations would require time-resolved imaging or perturbation experiments. Future work combining spatial transcriptomics with live-cell imaging, single-cell multi-omics, and larger cohorts will be important to further validate and refine these conclusions.

## Discussion

In this study, we used spatial transcriptomics to reconstruct the spatial and temporal organization of the cell cycle within human tumors. By integrating single-cell level cell-cycle inference with tissue context, we reveal how proliferative and quiescent states are arranged within the TME and how these states relate to tissue structure, vascularization, and cell-type composition. Our results demonstrate that the TME is not only a collection of diverse cell types, but also a dynamic mosaic of proliferative niches that reflect both local environmental constraints and intrinsic cellular programs.

A key finding of our work is that proliferating and non-proliferating cells form distinct spatial niches within tumors. Cycling cells, typically in G1, S, or G2/M phases, tend to cluster together in regions of higher cellular density of other cycling cells, while quiescent (G0) cells are enriched in regions of higher density of other quiescent cells. This suggests that proliferation is spatially coordinated, rather than randomly distributed across the tissue, and that microenvironmental factors such as oxygen availability, nutrient diffusion, or stromal composition may locally regulate the entry and exit from the cell cycle. The observation that endothelial and immune cells were predominantly quiescent across all tumors is consistent with their known roles in maintaining tissue stability and immune homeostasis. Endothelial quiescence, in particular, has been reported to suppress cancer invasiveness, whereas its loss leads to pathological angiogenesis and tumor expansion.

Our pseudotime analysis using SpaceCycle, a model based on the expectation–maximization algorithm, further extends this view by revealing oscillatory gene expression programs of cancer cells in situ. Unlike conventional discrete phase annotation, SpaceCycle infers a continuous proliferative trajectory, allowing estimation of phase durations and transition dynamics. Across tumor types, we observed distinct temporal signatures between cancer cell subpopulations, with some clusters exhibiting short G1 or G2/M phases, consistent with accelerated cell cycle progression or checkpoint loss, while others showed extended phases, indicative of arrest or stress responses. These differences point to intratumoral heterogeneity not only in static cell identity but also in proliferative behavior. Our study also demonstrates that incorporating dynamical inference into spatial omics can provide insights that are not accessible from static measurements alone. The integration of spatial coordinates with inferred cell cycle pseudotime bridges the gap between molecular dynamics and tissue architecture. This combination enables a “spatially resolved timeline” of proliferation, where neighboring cells can be placed along a continuum of activity. Such an approach can be extended beyond the cell cycle to study other cyclic or progressive biological processes, such as hypoxia adaptation, immune activation, therapy-induced senescence (TIS) or oncogene-induced senescence (OIS).

From a clinical and translational perspective, mapping proliferative organization within tumors has several implications. Spatially localized proliferation could serve as a biomarker for tumor aggressiveness, growth fronts, or therapy-resistant regions. Quiescent niches may contribute to therapeutic resistance by acting as sanctuaries for dormant cells that can later drive recurrence. Understanding how these regions are organized and regulated may inform new strategies to disrupt tumor architecture or reprogram the TME toward a less supportive state. Traditionally, the tumor proliferative index, which measures the relative fraction of actively dividing cells by quantifying the expression of a single marker, such as Ki-67 expression, PCNA or BrdU/Edu staining, is used for disease staging in cancer. Our approach combining spatial transcriptomics with quantitative modeling such as SpaceCycle not only allows to identify distinct cell cycle dynamics in subpopulations of cancer cells across the TME but also gives us gene expression dynamics in the cancer cells. This approach has the potential to revolutionize the cancer diagnosis and disease staging, as well as to identify novel targets for therapeutic development. In summary, our study presents a framework to study the spatial organization and dynamics of the cell cycle in tumors by combining discrete phase inference with a Fourier-based pseudotime model. We show that proliferation in tumors is spatially structured, that cycling and quiescent cells form distinct microenvironmental niches, and that dynamic modeling can uncover cell-type specific oscillatory programs underlying this organization. These findings highlight the importance of considering both space and time in understanding tumor growth and pave the way for spatially resolved dynamic modeling of other biological processes within complex tissues.

## Methods

### Clustering and cell type identification

We used the Python library Scanpy [50] (version 1.9.6) for preprocessing the data and for clustering analysis. We filtered cells that had fewer than 50 counts and genes that were expressed in fewer than 10 cells, normalized the counts for cell size effects and did `log1p` transform to make the data more comparable and less skewed, and suitable for clustering analysis. We did the principal components analysis, `n_comps=50` and obtained a *k−*NN graph for `n_neighbors=100`. Using this graph as the basis for Leiden clustering [20], we used the value of `resolution` that optimized the silhouette score [51] and the number of clusters, `resolution=0.5` for the melanoma sample, and `resolution=0.4` for breast and lung samples.

Tissue-specific cell type markers were obtained for cells that are normally present in the skin, breast, and lung from the CellMarker2.0 database [21]. For each Leiden cluster within the tissue we assigned the cell type based on the top 30 upregulated genes in the cluster. Based on the abundance of the gene markers, some clusters were identified to be composed of multiple cell types.

### Discrete cell-cycle phase identification

We obtained the cell cycle phase markers from Gene Ontology (GO) annotations, for identification of G0 phase, we looked at genes that were highly expressed in the quiescent subpopulation of human fibroblasts in from Riba et al [16]. We used the `scanpy.tl.score genes`, which is the Scanpy implementation of the approach introduced by Satija et al [52]. This gave us a score for the discrete cell cycle phases–G0, G1, S, G2/M–for every cell. We converted this score into a probability of the cell being in each phase

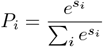

where *i* denotes the discrete cell cycle phases, G0, G1, S, G2/M, the, and *s*_*i*_ is the corresponding score. To improve the interpretability of our analysis, we created confusion matrix based on these probabilities and selected only those where we had a high degree of confidence in the phase identification (see supplementary Fig. S1)

### Comparison of distributions

Due to the large number of samples, the standard statistical tests like t-tests or Mann–Whitney U return extremely small p-values even for small, unimportant differences. We used Jensen-Shannon divergence [53], a symmetrized relative entropy measure based of Kull-back–Leibler divergence, to quantify how different the full distributions are without relying on significance testing.

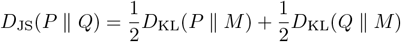

where *M* is the mixture distribution of *P* and 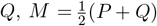, and *D*_KL_ is the Kullback–Leibler divergence,

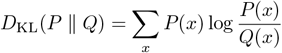

*D*_JS_ = 0 indicates that the two distributions are identical, *D*_JS_ = 1 indicates maximal possible divergence, achieved when the supports of *P* and *Q* are disjoint (i.e. samples can be perfectly assigned to one distribution or the other).

### Continuous cell-cycle phase identification using SpaceCycle

We assume that as the cells progress through the cell cycle, a subset of genes exhibit an oscillatory dynamics with respect to its cell cycle phase. We model this oscillatory dynamics using a Fourier series expansion,

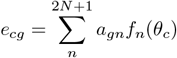

where, *e*_*cg*_ is the log-transformed gene expression of gene *g* in cell *c*; *f*_*n*_, the *n*^th^ component of the Fourier basis; *f* = (1 cos(2*πθ*_*c*_) · · · cos(2*Nπθ*_*c*_) sin(2*πθ*_*c*_) · · · sin(2*Nπθ*_*c*_))^⊤^; *a*_*gn*_, the *n*^th^ Fourier coefficient of gene *g*; and, *θ*_*c*_, the continuous cell cycle phase of cell *c*.

We implemented an expectation-maximization (EM) algorithm, an iterative method to find (local) maximum likelihood, to estimate the Fourier coefficients *a*_*gn*_ (the model parameters), and the cell cycle phase *θ*_*c*_ (the latent variable). This approach is similar to a method previously used to study the circadian clock in humans using bulk RNA-seq samples [54]. Under the independent and identically distributed assumption, the likelihood can be written as

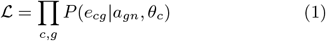

We assume that the log-transformed gene expression follows a Normal distribution,

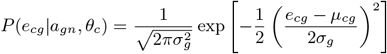

where *µ*_*cg*_ = Σ_*n*_ *a*_*gn*_*f*_*n*_(*θ*_*c*_), is the predicted gene expression of gene *g* in cell *c*

#### E step

In the E-step we obtain the expected value of the log-likelihood with respect to the conditional probability of a cell having a phase *θ*_*c*_ given that the gene expression data, *e*_*cg*_ and the Fourier coefficients *a*_*gn*_,

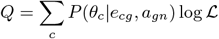

where,

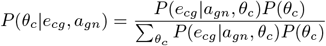

#### M step

In the M-step we maximize the expected likelihood with respect to the Fourier coefficients

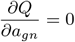

This gives us,

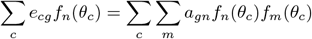

which can be expressed as a matrix equation,

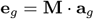

Thus we can obtain the Fourier coefficients for the genes,

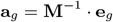

### Identification of phase transitions

SpaceCycle assigns each cell a continuous and periodic phase value, *θ*, based on its transcription profile. However, these raw *θ* values do not directly correspond to biological cell-cycle phases. To align *θ* with biological progression, we first require that *θ* increases from 0 to 1 in the order G1 → S → G2/M. We binned *θ* into equal-sized intervals and calculated, for each bin, the proportions of cells annotated as G1, S, and G2/M using the discrete phase scores (see Supplementary Fig. S4). The correct orientation of *θ* was identified by checking that the peak proportion of G1 precedes S, and S precedes G2/M. If the order was reversed, we transformed the phases by *θ* → 1 − *θ*. Once the correct orientation was set, we estimated the boundaries between G1/S, S/G2M, and G2M by scanning across all possible triplets of boundary positions. For each combination, we compared the resulting phase assignments with the discrete phase labels and selected the boundaries that maximized the agreement with the observed proportions of G1, S, and G2/M cells.

## Resources

### Code

https://github.com/MolinaLab-IGBMC/SpaceCycle

### Processed data

https://zenodo.org/records/17749853

## Acknowledgments

This work was supported by the Chan Zuckerberg Initiative (CZI) Single-Cell Biology Data Insights program through the project *Space-time Regulation of Biological Cycles in Cancer (DI3) to decipher circadian and cell cycle dynamics in cancer via context-dependent periodic manifold modeling: from regulation to new opportunities for chrono-treatments*.

GZ is supported by a PhD fellowship from La Ligue Nationale Contre le Cancer.

This work of the Interdisciplinary Thematic Institute IMCBio+, as part of the ITI 2021–2028 program of the University of Strasbourg, CNRS, and INSERM, was supported by IdEx Unistra (ANR-10-IDEX-0002) and by the SFRI-STRAT’US project (ANR-20-SFRI-0012) and EUR IMCBio (ANR-17-EURE-0023) under the framework of the France 2030 Program.

## Supplemental information

**Figure S1.**
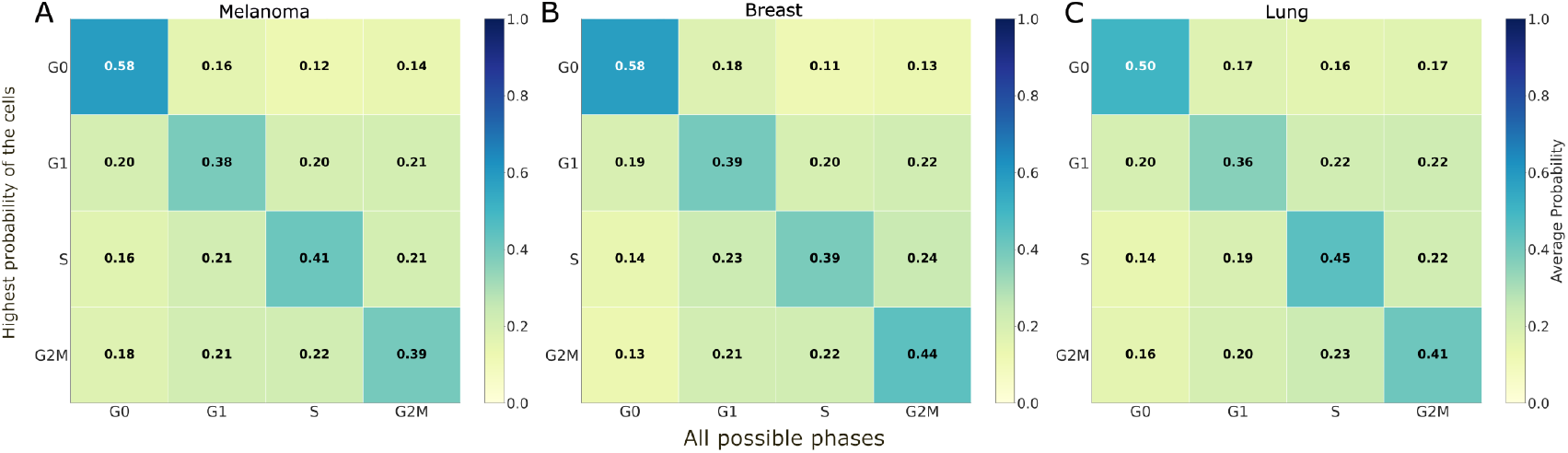
Confusion matrix between maximum probability of cells being in a one of the cell cycle phases and the individual probabilities of cell being in one of the cell cycle phases.

**Figure S2.**
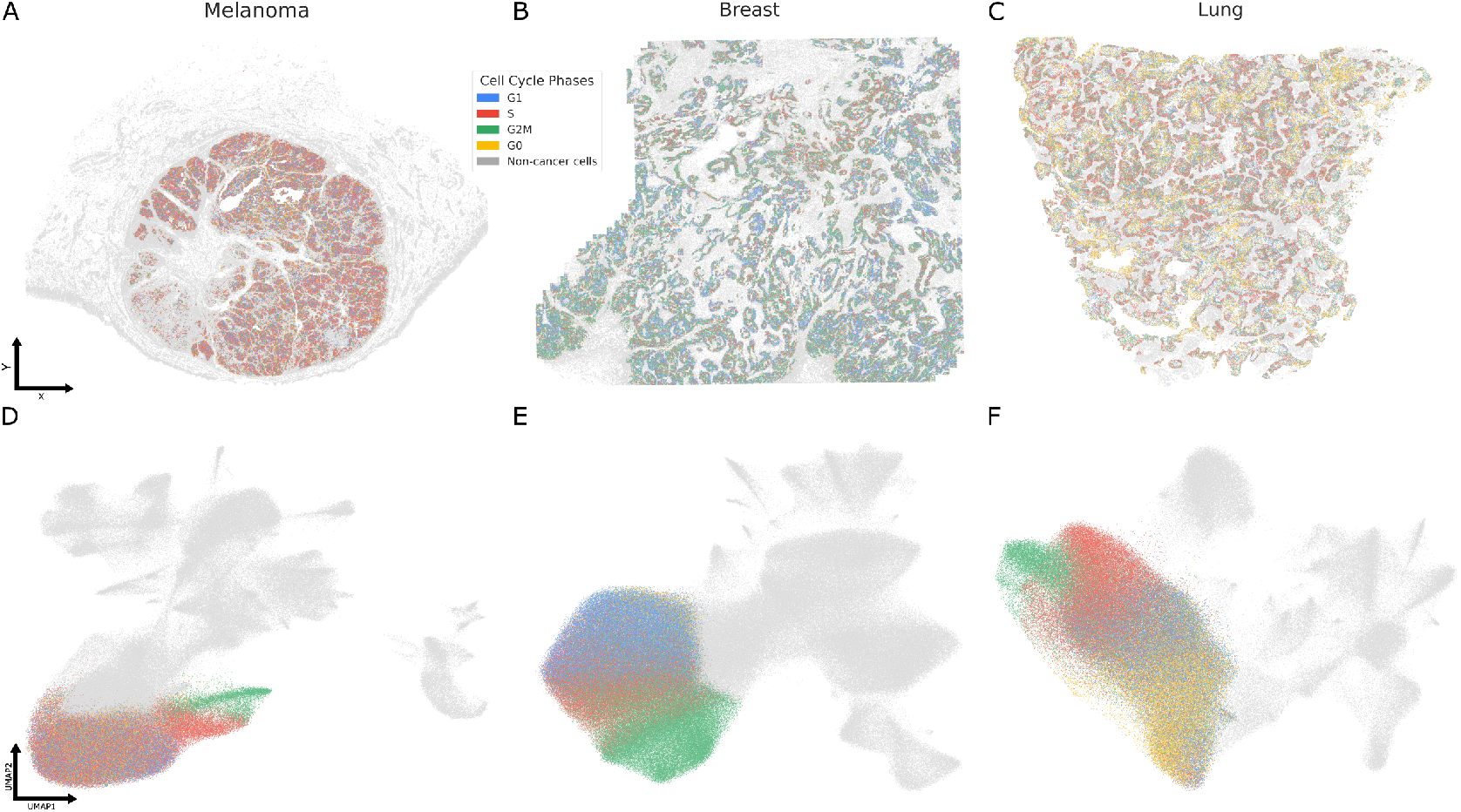
Discrete cell cycle phase of cancer cells. **A-C:** Spatial map discrete cell cycle phases–G0 (in yellow), G1 (in blue), S (in red), and G2/M (in green)–of cancer cells in the melanoma, breast and lung tumors. **D-F:** Corresponding cells visualized on the UMAP embedding. Grey denote the non-cancerous cells.

**Figure S3.**
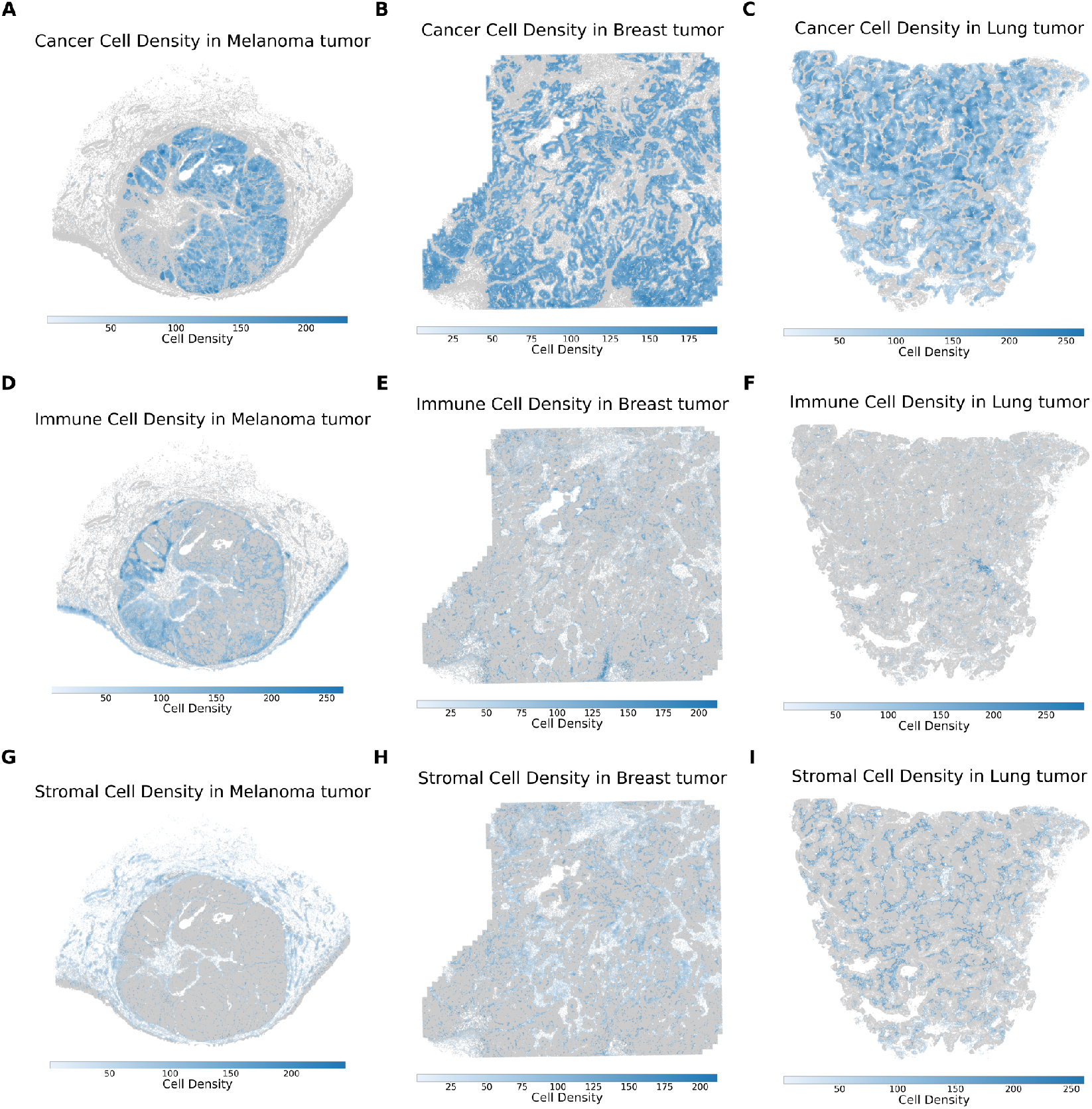
Cellular densities by cell types. Cellular densities of major cell types, **A-C** cancer cells, **D-F** immune cells, **G-I** stromal cells in the melanoma, breast, and lung sample

**Figure S4.**
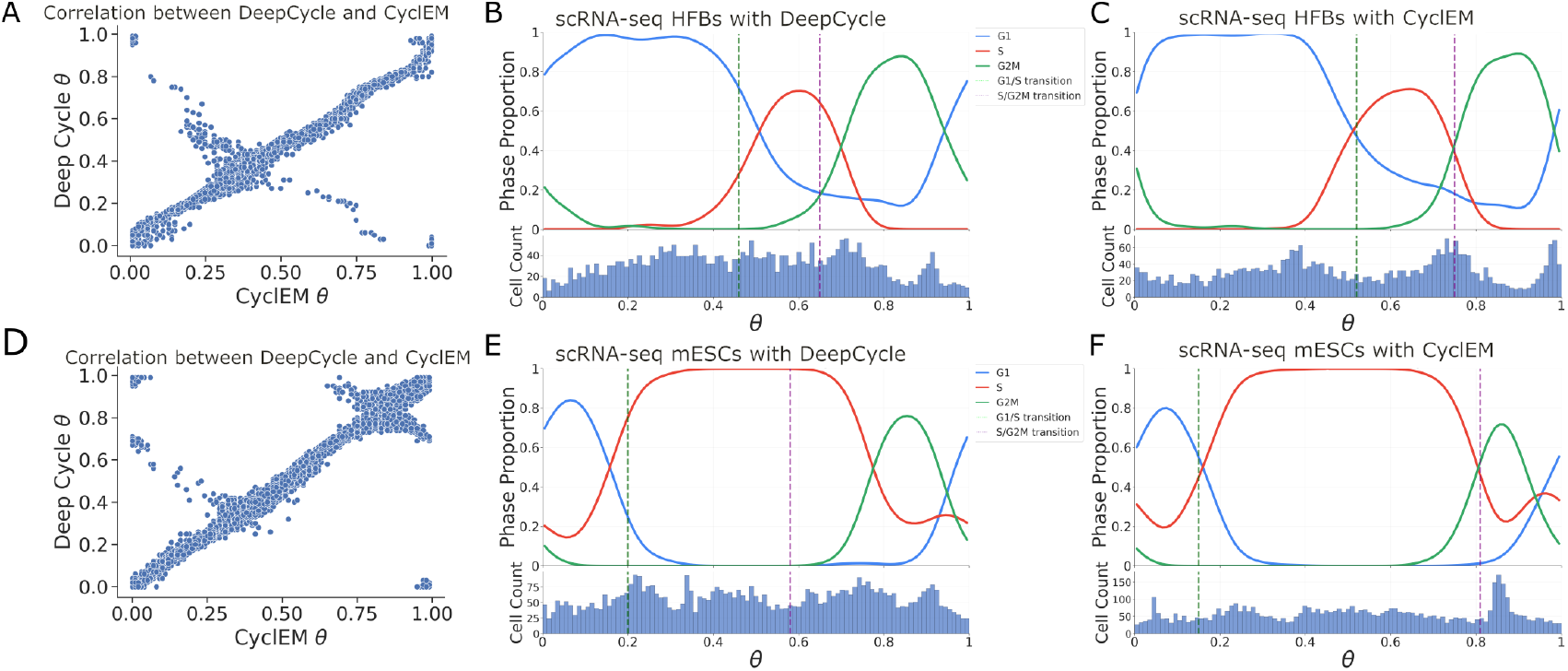
Validation of SpaceCycle with single-cell RNA-seq data. **A:** Scatterplot showing comparison of continuous cell cycle phase, *θ*, obtained with DeepCycle, *x*-axis, versus SpaceCycle, on *y*-axis for single-cell RNA-seq data of human fibroblasts from Riba et al [16]. **B** phase inference using DeepCycle, bottom panel: histogram showing distribution of cell cycle phases when binned into 100 bins, top panel: proportions of cells in G1, S and G2/M in each bin, discrete phase identification was done as described in the Methods section, the dotted line represent the G1/S (in green) and the S/G2M (in purple) transitions respectively. **C:** Corresponding plots when phase inference was performed using CyclEM. **D, E** and **F:** corresponding plots for scRNA-seq from mouse embryonic stem cells from Riba et al [16].

**Figure S5.**
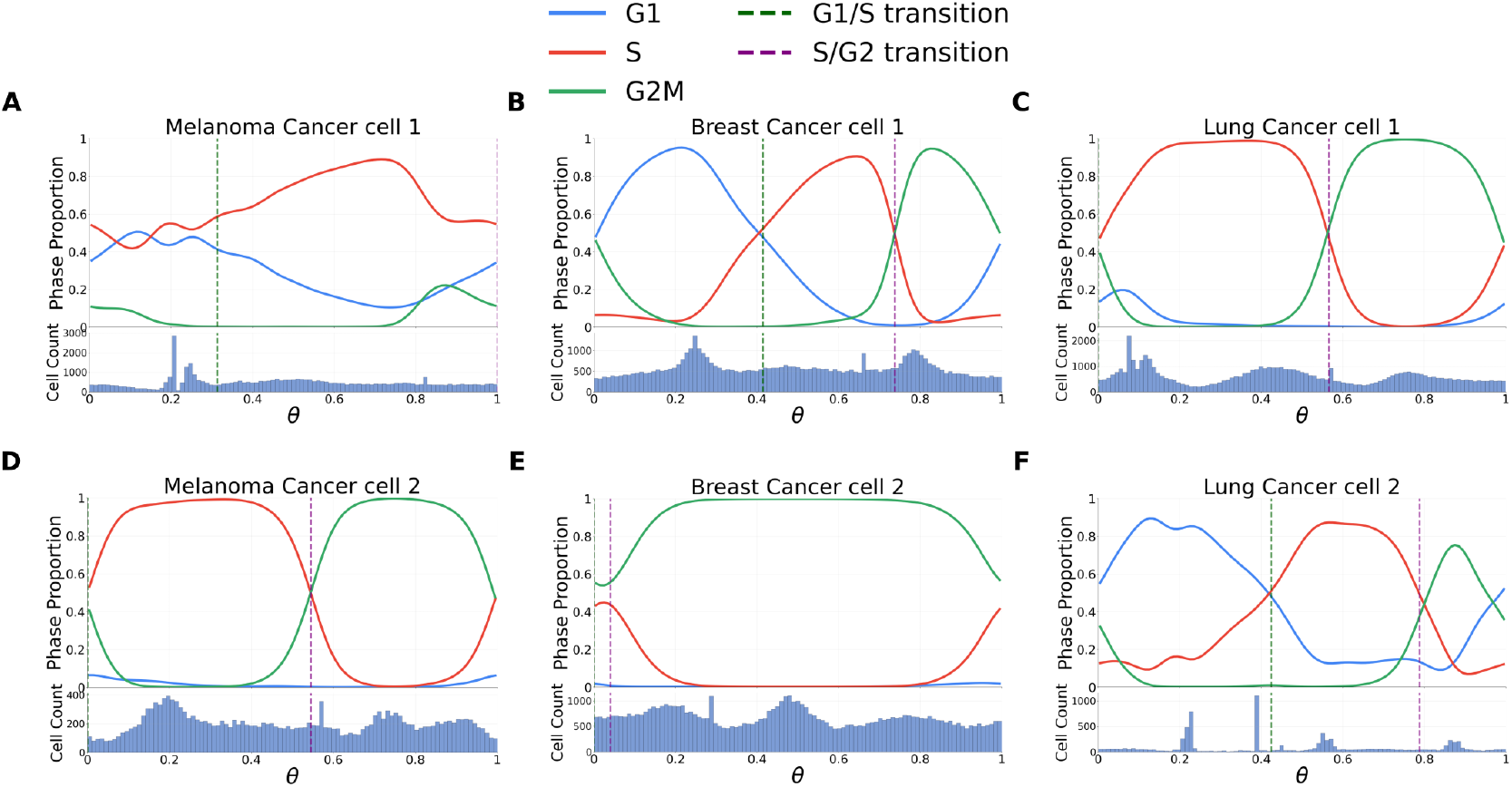
Continuous phase distributions and phase transitions in cell clusters. Distributions of continuous phases, proportions of cells in discrete phases, and phase transition boundaries for clusters in melanoma, cancer cell 1 in **A** and cancer cell 2 in **D**, breast, cancer cell 1 in **B** and cancer cell 2 in **E**, and lung, cancer cell 1 in **C** and cancer cell 2 in **D**

**Table S1.** Tumor sample general features and quality metrics.

| Tumor sample | Number of cells | Number of transcripts | RIN | DV200 |
| --- | --- | --- | --- | --- |
| Melanoma | 468,138 | 160,181,929 | 1.9 | 60.73 |
| Breast | 713,121 | 490,398,542 | 2.7 | 87.37 |
| Lung | 353,762 | 144,388,044 | 4.3 | 66.97 |

**Table S2.** Top 4 cell types by population in each tumor, with the number of cells.

| Tumor sample | Cell type | Number of cells |
| --- | --- | --- |
| Melanoma | Cancer cells | 179,905 |
| Melanoma | Endothelial cells and pericytes | 90,511 |
| Melanoma | Macrophages | 79,726 |
| Melanoma | Monocytes | 32,994 |
| Breast | Cancer cells | 305,652 |
| Breast | Macrophages | 95,632 |
| Breast | Cancer associated fibroblasts | 88,098 |
| Breast | Cancer stem cells | 70,921 |
| Lung | Cancer cells | 209,047 |
| Lung | Secretory cells | 33,064 |
| Lung | T cells | 21,596 |
| Lung | Macrophages | 20,770 |

**Table S3.** Genes used for discrete cell cycle phase identification.

| Phase | Gene markers |
| --- | --- |
| G0 | COL1A, LRP1, CDKN1A, CD248 |
| G1 | CCND1, CDK4, CDK6, CDKN1B, RB1, TP53 |
| S | DNMT1, MCM2, MCM6, MSH2, MSH3, MSH6, PCNA |
| G2M | APC, AURKA, AURKB, BIRC5, BUB1, CCNB1, ESCO2, FOXM1, MKI67, MYBL2, PLK1 |

## References

1 J. Folkman, “Tumor angiogenesis: therapeutic implications”, New England Journal of Medicine 285 (1971).

2 D. Hanahan and J. Folkman, “Patterns and emerging mechanisms of the angiogenic switch during tumorigenesis”, Cell 86 (1996).

3 K. J. Kim et al., “Inhibition of vascular endothelial growth factor-induced angiogenesis suppresses tumour growth in vivo”, Nature 362 (1993).

4 N. A. Bhowmick et al., “Stromal fibroblasts in cancer initiation and progression”, Nature 432 (2004).

5 A. Orimo et al., “Stromal fibroblasts present in invasive human breast carcinomas promote tumor growth and angiogenesis through elevated sdf-1/cxcl12 secretion”, Cell 121 (2005).

6 R. Kalluri and M. Zeisberg, “Fibroblasts in cancer”, en, Nat. Rev. Cancer 6 (2006).

7 S. L. Topalian et al., “Immune checkpoint blockade: a common denominator approach to cancer therapy”, Cancer Cell 27 (2015).

8 D. M. Pardoll, “The blockade of immune checkpoints in cancer immunotherapy”, Nature Reviews Cancer 12 (2012).

9 P. Sharma et al., “Immune checkpoint therapy—current perspectives and future directions”, Cell 186 (2023).

10 K. H. Chen et al., “RNA imaging. spatially resolved, highly multiplexed RNA profiling in single cells”, en, Science 348 (2015).

11 J.-R. Lin et al., “Highly multiplexed imaging of single cells using a high-throughput cyclic immunofluorescence method”, Nature Communications 6 (2015).

12 J. Liu et al., “Concordance of merfish spatial transcriptomics with bulk and single-cell rna sequencing”, Life Science Alliance 6 (2022).

13 S. P. Castillo et al., “The tumour ecology of quiescence: niches across scales of complexity”, Seminars in Cancer Biology 92 (2023).

14 P. Baldominos et al., “Quiescent cancer cells resist t cell attack by forming an immunosuppressive niche”, Cell 185 (2022).

15 G. La Manno et al., “RNA velocity of single cells”, en, Nature 560 (2018).

16 A. Riba et al., “Cell cycle gene regulation dynamics revealed by RNA velocity and deep-learning”, en, Nat. Commun. 13 (2022).

17 M. K. Nariya et al., “Single-cell multiomics reveals the oscillatory dynamics of mrna metabolism and chromatin accessibility during the cell cycle”, Cell Reports 44 (2025).

18 A. R. Lederer et al., “Statistical inference with a manifold-constrained rna velocity model uncovers cell cycle speed modulations”, Nature Methods 21 (2024).

19 Vizgen, Merfish ffpe human immuno-oncology data set, 2022.

20 V. A. Traag et al., “From louvain to leiden: guaranteeing well-connected communities”, en, Sci. Rep. 9 (2019).

21 C. Hu et al., “CellMarker 2.0: an updated database of manually curated cell markers in human/mouse and web tools based on scRNA-seq data”, en, Nucleic Acids Res. 51 (2023).

22 G. Palla et al., “Squidpy: a scalable framework for spatial omics analysis”, en, Nat. Methods 19 (2022).

23 I. Tirosh et al., “Dissecting the multicellular ecosystem of metastatic melanoma by single-cell RNA-seq”, en, Science 352 (2016).

24 B.-Z. Qian and J. W. Pollard, “Macrophage diversity enhances tumor progression and metastasis”, en, Cell 141 (2010).

25 R. A. Franklin et al., “The cellular and molecular origin of tumor-associated macrophages”, en, Science 344 (2014).

26 E. Sahai et al., “A framework for advancing our understanding of cancer-associated fibroblasts”, en, Nat. Rev. Cancer 20 (2020).

27 S. Liu et al., “Breast cancer stem cells transition between epithelial and mesenchymal states reflective of their normal counterparts”, en, Stem Cell Reports 2 (2014).

28 P. C. Boutros et al., “Prognostic gene signatures for non-small-cell lung cancer”, Proceedings of the National Academy of Sciences 106 (2009).

29 M. L. Forsythe et al., “Molecular profiling of non-small cell lung cancer”, PLOS ONE 15, edited by R. A. De Mello (2020).

30 R. Nussinov et al., “Molecular principles underlying aggressive cancers”, Signal Transduction and Targeted Therapy 10 (2025).

31 J. W. Franses et al., “Stromal endothelial cells directly influence cancer progression”, en, Sci. Transl. Med. 3 (2011).

32 P. Leone et al., “Endothelial cells in tumor microenvironment: insights and perspectives”, Frontiers in Immunology 15 (2024).

33 Z. Wang et al., “Cancer-associated fibroblasts suppress cancer development: the other side of the coin”, Frontiers in Cell and Developmental Biology 9 (2021).

34 D. Yang et al., “Cancer-associated fibroblasts: from basic science to anticancer therapy”, Experimental & Molecular Medicine 55 (2023).

35 Y. Miyai et al., “Cancer-associated fibroblasts that restrain cancer progression: hypotheses and perspectives”, Cancer Science 111 (2020).

36 D. Ackerman and M. C. Simon, “Hypoxia, lipids, and cancer: surviving the harsh tumor microenvironment”, Trends in Cell Biology 24 (2014).

37 M. Esteves et al., “The effects of vascularization on tumor development: a systematic review and metaanalysis of pre-clinical studies”, Critical Reviews in Oncology/Hematology 159 (2021).

38 A. Karsch-Bluman and O. Benny, “Necrosis in the tumor microenvironment and its role in cancer recurrence”, in Tumor microenvironment (Springer International Publishing, 2020).

39 A. R. Barr and F. Gergely, “Aurora-A: the maker and breaker of spindle poles”, en, J. Cell Sci. 120 (2007).

40 T. Courtheoux et al., “Aurora a kinase activity is required to maintain an active spindle assembly checkpoint during prometaphase”, en, J. Cell Sci. 131 (2018).

41 J. den Hollander et al., “Aurora kinases a and B are up-regulated by myc and are essential for maintenance of the malignant state”, en, Blood 116 (2010).

42 N. A. Borah and M. M. Reddy, “Aurora kinase B inhibition: a potential therapeutic strategy for cancer”, en, Molecules 26 (2021).

43 M. A. Edelbrock et al., “Structural, molecular and cellular functions of MSH2 and MSH6 during DNA mismatch repair, damage signaling and other noncanonical activities”, en, Mutat. Res. 743–744 (2013).

44 M. H. Theus et al., “EphB3 limits the expansion of neural progenitor cells in the subventricular zone by regulating p53 during homeostasis and following traumatic brain injury”, en, Stem Cells 28 (2010).

45 J. Holmberg et al., “EphB receptors coordinate migration and proliferation in the intestinal stem cell niche”, en, Cell 125 (2006).

46 Z. Xiao et al., “EphB3 receptor suppressor invasion, migration and proliferation in glioma by inhibiting EGFR-PI3K/AKT signaling pathway”, en, Brain Res. 1830 (2024).

47 J.-J. Li et al., “EphB3 stimulates cell migration and metastasis in a kinase-dependent manner through Vav2-Rho GTPase axis in papillary thyroid cancer”, en, J. Biol. Chem. 292 (2017).

48 K. Harikrishnan et al., “Cell derived matrix fibulin-1 associates with epidermal growth factor receptor to inhibit its activation, localization and function in lung cancer calu-1 cells”, en, Front. Cell Dev. Biol. 8 (2020).

49 T. S. Mok et al., “Gefitinib or carboplatin-paclitaxel in pulmonary adenocarcinoma”, en, N. Engl. J. Med. 361 (2009).

50 F. A. Wolf et al., “Scanpy: large-scale single-cell gene expression data analysis”, Genome Biology 19 (2018).

51 P. J. Rousseeuw, “Silhouettes: a graphical aid to the interpretation and validation of cluster analysis”, Journal of Computational and Applied Mathematics 20 (1987).

52 R. Satija et al., “Spatial reconstruction of single-cell gene expression data”, en, Nat. Biotechnol. 33 (2015).

53 D. Endres and J. Schindelin, “A new metric for probability distributions”, IEEE Transactions on Information Theory 49 (2003).

54 L. Talamanca et al., “Sex-dimorphic and age-dependent organization of 24-hour gene expression rhythms in humans”, Science 379 (2023).

